# Recombineering in *Vibrio natriegens*

**DOI:** 10.1101/130088

**Authors:** Henry H. Lee, Nili Ostrov, Michaela A. Gold, George M. Church

## Abstract

Here, we show that λ-Red homologs found in the *Vibrio*-associated SXT mobile element potentiate allelic exchange in *V. natriegens* by ~10,000-fold. Specifically, we show SXT-Beta (s065), SXT-Exo (s066), and λ-Gam proteins are sufficient to enable recombination of single- and double-stranded DNA with episomal and genomic loci. We characterize and optimize episomal oligonucleotide-mediated recombineering and demonstrate recombineering at genomic loci. We further show targeted genomic deletion of the extracellular nuclease gene *dns* using a double-stranded DNA cassette. Continued development of this recombination technology will advance high-throughput and large-scale genetic engineering efforts to domesticate *V. natriegens* and to investigate its rapid growth rate.

## Main text

We have previously proposed the fast growing marine bacterium *Vibrio natriegens* as a powerful bacterial host and reported foundational genomics resources for its utilization (*1*). While methods for genome modification of *V. natriegens* by homologous recombination have recently been reported, they are laborious and protract experimental time. Plasmid-based integration methods require extensive cloning, conjugation, and a strong negative selection for elimination of the plasmid backbone (*2*). Similarly, recombination of double-stranded DNA cassettes by natural competence, though attractive due to its efficiency, requires cloning of unwieldy homology arms up to 3 kb and extended incubation times (*3*). Development of one-step recombineering method which tolerates short homology arms, particularly with oligonucleotides, would be an attractive advancement for genomic manipulation of *V. natriegens*.

Recombineering is a powerful method for precise DNA editing, enabling *in vivo* construction of mutant alleles and structural changes such as insertions and deletion of genes (*4–9*). These mutations can be introduced by allelic exchange between the target sequence and recombinant single- or double-stranded DNA, potentiated by expression of powerful homologous recombination (HR) proteins found in bacteriophages (*5*). λ-Beta, the most well-studied phage recombinase, has been shown to enhance HR in *E. coli* by ~10,000-fold over the basal mutation rate (*10*). Unfortunately, λ-phage Red Beta does not sustain this efficiency in diverse bacteria (*10–17*). In addition, recombinant expression of some λ-phage Red Beta homologues in *E. coli* displays attenuated HR activity, suggesting potential reliance on host-specific machinery (*18, 19*). To maximize the chances of finding a functional recombinase in *V. natriegens*, we searched for λ-phage Red homologs that had been previously identified in *Vibrio* species.

Here, we demonstrate the use of SXT recombinase proteins in *V. natriegens* for targeted mutagenesis using single- and double-stranded DNA. The SXT mobile genetic element, originally isolated from an epidemic strain of *Vibrio cholerae*, encodes homologous proteins to *E. coli* ssb (s064, single-strand binding protein), λ-Beta (s065, single-strand annealing protein), and λ-Exo (s066, alkaline exonuclease protein) (*18, 20, 21*). SXT-Beta shares 43.6% identity with λ-Beta protein, whereas SXT-Exo shares only 25.1% identify with its λ homolog (Fig. 1a,b). We were unable to find *Vibrio*-specific homologs of λ-Gam. Importantly, previous studies have shown that in *E. coli*, SXT-Beta enhances oligo HR to levels comparable to λ-Beta and RecT, whereas SXT-Beta and SXT-Exo together displays 50-fold lower dsDNA HR activity compared to λ-Beta-Exo and RecET (*18, 19*). We thus synthesized SXT-Beta-Exo *de novo* and placed it in an operon with λ-Gam for recombinant expression in *V. natriegens* (Methods).

**Figure 1.**
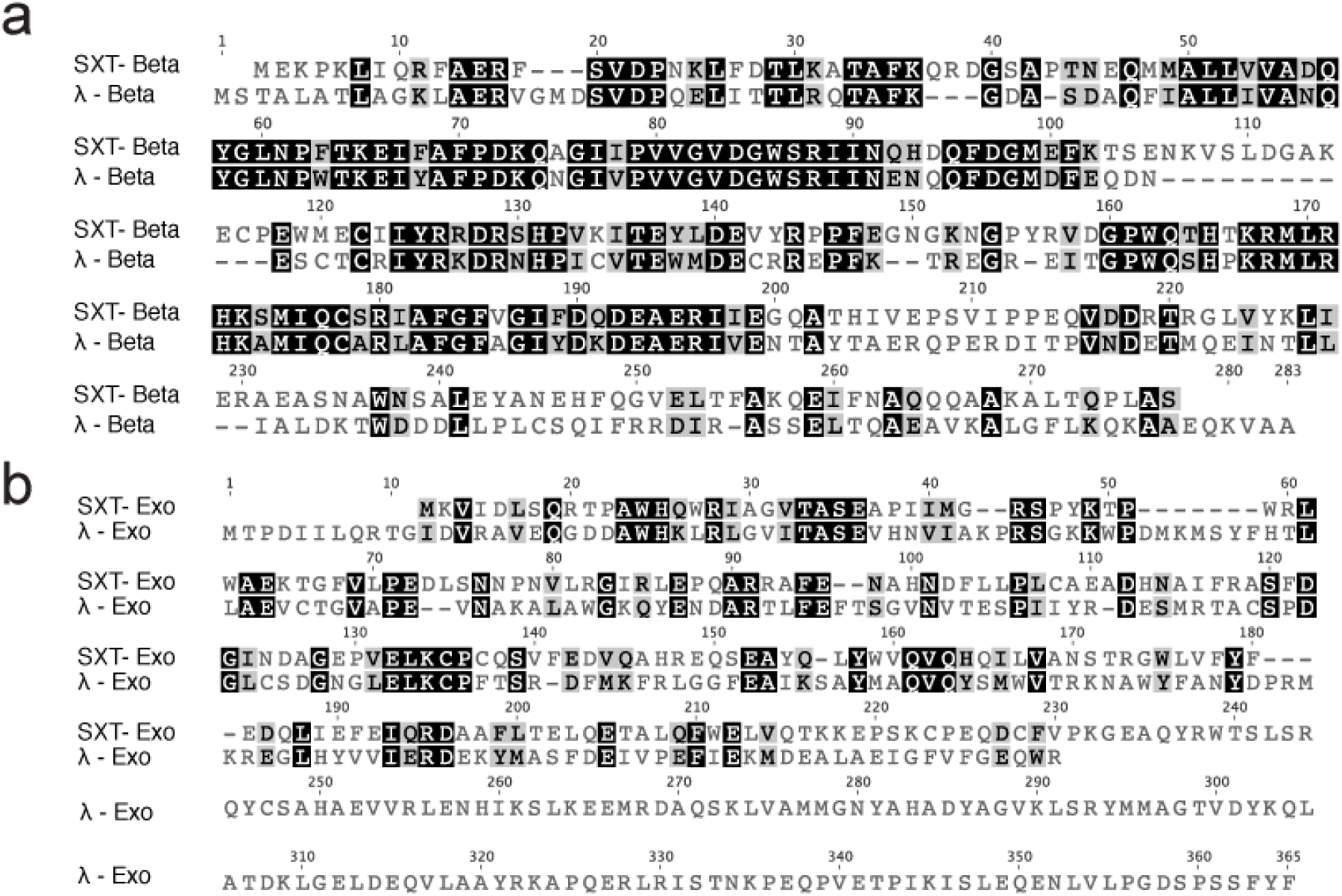
Local alignment of SXT-Beta and SXT-Exo with λ-Beta and λ-Exo. Alignments were performed as Global alignment with free-end gaps (Geneious) with BLOSUM62, gap open penalty 12 and gap extension penalty 3. (a) SXT-Beta and λ-Beta are 43.6% identical and 58.5% similar. (b) SXT-Exo and λ-Exo are 25.1% identical and 38.4% similar.

To quantify recombination events, we designed a selection-based assay where a recombinogenic oligonucleotide rescues an otherwise inactive antibiotic marker. Specifically, we introduced a premature stop codon to the spectinomycin gene by mutating a single base, and cloned it onto a kanamycin-resistant pRST vector (Methods). Successful recombinants would result in colony formation on spectinomycin plates. We first assessed the background spectinomycin reversion rate with or without oligonucleotide. We found the spontaneous reversion rate to be < 10^-10^ cells in the absence of any recombination machinery (no spectinomycin resistant colonies could be detected). Importantly, we found that upon introduction of oligonucleotide, strains expressing SXT recombination proteins yielded ~30-fold the number of colonies compared to no oligo control, and 100-fold the number of colonies compared to strain lacking any recombinase (Fig. 2a). Notably, we found λ-Beta to be > 2-fold less efficient than SXT-Beta; further work is required to better assess this difference (Methods).

**Figure 2.**
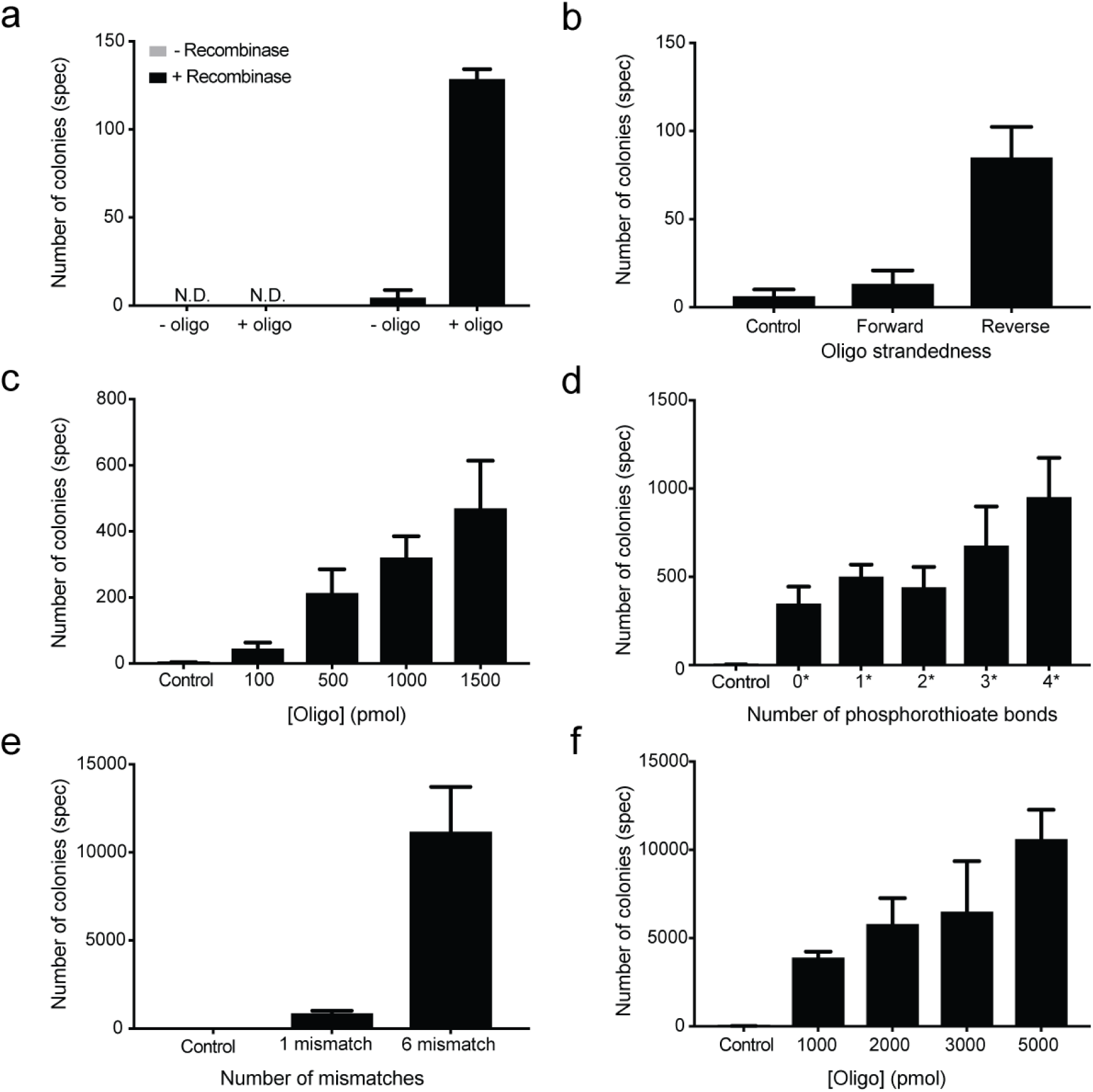
Optimization of SXT-mediated oligonucleotide recombination. A oligonucleotide carrying a reversion was used to revert a premature stop codon in an episomal spectinomycin gene. Recombinants expressed as number of spectinomycin-resistant colonies. Experiments performed by lac-inducible expression of an operon consisting of SXT-beta, SXT-exo, and λ-Gam. Oligonucleotide are 90-mers. N.D. is Not Detected. (a) Spontaneous mutation or SXT-mediated recombination (fixspec_R; 250 pmol). (b) Oligonucleotide targeting the leading or lagging strand (fixspec_F, fixspec_R; 250 pmol). (c) Titration of oligonucleotide (fixspec_R). (d) Number of 5’ terminal phosphorothioate bonds (oligos BS_rP0 - BS_rP4, 1500 pmol). (e) Number of consecutive mismatches in oligonucleotide (fixspec_R, fixSpec_6MM_R; 1500 pmol). (f) Titration of 6-mismatch oligonucleotide (fixSpec_6MM_R).

We next proceeded to improving SXT-mediated recombination efficiency by optimizing induction times and oligonucleotide properties such as concentration, strandedness, and number of phosphorothioate bonds. We first explored conditions for sufficient induction of recombination proteins. Whereas standard *E. coli* recombineering protocols utilize short inductions, we found that overnight induction yielded 3-fold more recombinants than short induction upon subculturing (*5, 22*) (Data not shown).

Previous studies have shown altered efficiency of λ-Beta recombination in *E.coli* based on the target’s strandedness, as the mechanism of incorporation occurs via the creation of an Okazaki-like primer during chromosome replication (*23*). As a result, oligonucleotides targeting the lagging strand are more recombinogenic than those targeting the leading strand. We tested for this hallmark of Beta-like recombination using complementary oligonucleotides, and found the “forward” oligonucleotide to be 6-fold less recombinogenic than the “reverse” oligonucleotide (Fig. 2b). Based on this observation, we performed subsequent plasmid recombination assays with oligonucleotides targeting the “reverse” strand.

We next sought to determine the required oligonucleotide concentration for optimal recombination efficiencies. We observed that recombination efficiency increased linearly with increasing oligonucleotide concentration (Fig. 2c). Notably, the highest tested concentration of oligonucleotides, equivalent to 90,300 oligonucleotide molecules per cell (1500 picomoles for ~10^10^ electrocompetent cells), have not yet reached saturation of recombination efficiency. Considering a multi-copy plasmid target, this high ratio of oligos per cell strongly suggests that oligonucleotides are not efficiently recombined and are likely subject to degradation.

To probe potential sources of oligonucleotide degradation, we first modified our oligonucleotides with 5’ phosphorothioate bonds. Increased number of 5’ phosphorothioate bonds has been shown to increase recombination efficiency by preventing nonspecific exonuclease activity (*5, 24, 25*). We found mild enhancement of recombination efficiency with addition of 1, 2, or 3 phosphorothioates, reaching 2-fold improvement using 4 phosphorothioates, similar to previous reports for recombination by λ-Beta in *E. coli* (*5*) (Fig. 2d).

Another form of oligonucleotide degradation is the repair of DNA mismatches by the endogenous mismatch repair (MMR) system (*26*). Previous studies have found that oligonucleotides carrying 6 mismatches effectively evade the endogenous *E. coli* MMR system (*27, 28*). We introduced 5 additional mismatched bases while maintaining synonymous codons. We found oligonucleotide with 6 mismatches to be ~13-fold more recombinogenic than the one with a single mismatch (Fig. 2e). Finally, we increased the concentration this oligonucleotide to 5000 pmoles, equivalent to 301,000 strands per electrocompetent cell, which yielded more than 10^4^ recombinants (Fig. 2f). Taken together, our results show that SXT-Beta potentiates oligonucleotide-mediated HR to an episomal target in *V. natriegens* by ~10,000-fold.

To assess oligonucleotide-mediated allelic exchange with genomic loci, we sought a chromosomal target which can be used as a selection when mutagenized. We chose 5-fluoroorotic acid (FOA) counterselection, which was originally established in *S. cerevisiae* and has been adapted for use in some Gram-negative bacteria (*29, 30*). FOA counterselection depends on the conversion of the uracil analog, 5-FOA, into a highly toxic compound by pyrimidine metabolic enzyme Orotidine-5'-phosphate decarboxylase, encoded by *S. cerevisiae* URA3. Cells with a functional URA3 gene are sensitive to 5-FOA, while cells with inactive URA3 survive. For wild-type *V. natriegens*, we found that colonies will form over time despite high levels of 5-FOA, necessitating replica plating to improve selection (Methods).

We targeted the *pyrF* (chr1), the *V. natriegens* URA3 homolog, for introduction of a premature stop codon by oligonucleotide-mediated recombination. Following electroporation, colonies were grown, replica plated on 5-FOA plates, and the target loci was sequenced by Sanger (Methods). We found 2 of 9 colonies which carried the desired *pyrF* mutation with the remaining colonies showing aberrant deletions, indicating that FOA induces unwanted mutagenesis. Thus, we next sought to directly quantify oligonucleotide-mediated genomic recombination by deep sequencing.

We targeted four genes for introduction of in-frame stop codons: *dns* (chr1), *mutS* (chr1), *lacZ* (chr1), and *xds* (chr2). To evade endogenous MMR, mutations were designed such that 6 contiguous mismatches would result in 2 consecutive premature stops codons (*27, 28*). We pooled and electroporated oligonucleotides into cells expressing SXT recombination proteins, then quantified the abundance of each allele by deep sequencing (Methods). In the absence of selection, we were able to detect allelic exchange for the *xds* target 1 in ~10^6^ reads whereas we were unable to observe mutations at the other 3 targets at our sequencing depth (*mutS,* ~10^5^ reads; *dns and lacZ,* ~10^6^ reads). Performing a second round of recombineering, which increases the penetrance of allelic exchange in *E. coli*, did not improve the detection of these recombination events (*5*). Our initial optimization of SXT-mediated recombineering enables direct genomic editing in *V. natriegens* and further work is ongoing to improve its efficiencies to rival that of λ-Red recombineering in *E.coli* (*10*).

Interestingly, we found the genomic copy number of *xds* to be ~4-fold less than the other 3 targets, indicating *xds* is further from the origin of replication (Methods) Given the fewer *xds* targets relative to the other loci and the rapid generation time of *V. natriegens*, it is interesting to consider potential host-specific processes that may hinder techniques for genome modifications. Further work to pinpoint the underlying molecular mechanisms that impede genomic recombination will be required to enable selection-free recombineering.

In addition to oligonucleotides, recombineering can be achieved by double-stranded cassettes which contain regions of homology to the target loci. It has been previously shown that λ-Exo and λ-Gam are required in addition to λ-Bet protein for dsDNA recombineering in *E. coli (31)*. Similarly, we show that co-expression of SXT-Exo and λ-Gam in addition to SXT-Beta is sufficient for dsDNA recombineering in *V. natriegens*.

We chose to delete *dns*, one of two extracellular DNAse genes in *V. natriegens*, which is homologous to *endA* in *E. coli.* Strains of *E. coli* carrying *endA1* allele are deficient in DNAse activity and are useful as cloning and sequencing strains (*32*). Recently, a *Vibrio natriegens* Δ*dns* strain was shown to improve plasmid yield (*2*). We constructed a double-stranded DNA cassette by assembly PCR which consisted of the spectinomycin antibiotic gene flanked on both ends by 500 bp genomic sequences immediately downstream and upstream of *dns*. Cassettes were PCR amplified with primers containing zero or two phosphorothioates at the 5’ end, yielding 4 separate cassette types. We electroporated 1 μg (~750 fmol) of each cassette into *V. natriegens* strain expressing the SXT recombination operon. Resulting spectinomycin resistant colonies were screened by PCR, as recombination would yield a 2.1 kb product rather than 1.7 kb for wild type (Fig. 3a). Notably, we only obtained colonies when the input cassette contained 5’ phosphorothioates on at least one strand, indicating that evasion of endogenous nucleases is critical in *V. natriegens*. We found 1 of 8 transformants to yield the 2.1 kb PCR product and we confirmed this putative *dns* deletion mutant by whole genome sequencing (Fig. 3b).

**Figure 3.**
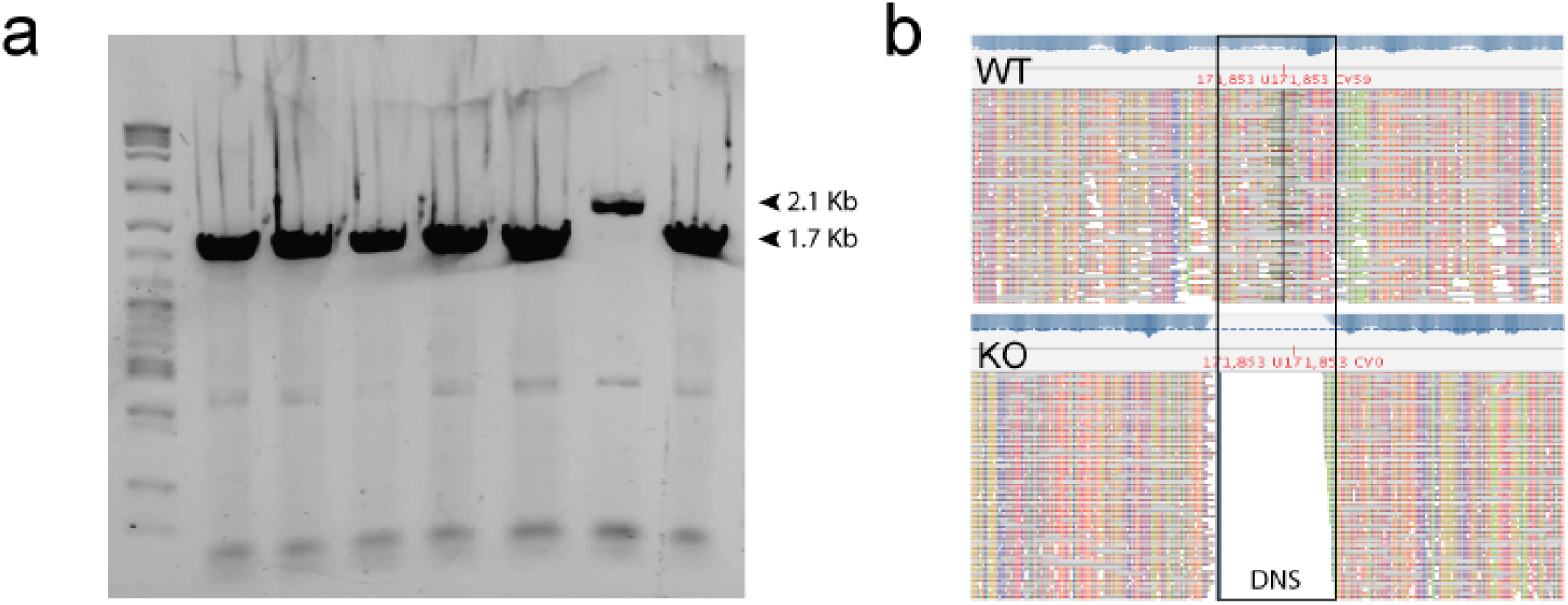
Targeted deletion of *dns* by SXT-mediated dsDNA recombination. (a) Putative dsDNA recombinants were screened by PCR with primers surrounding the *dns* locus. Successful recombination revents result in replacement of *dns* with a spectinomycin resistance marker. Wild type locus yields a 1.7 kb band whereas the recombinant locus yields a 2.1 kb band which includes spectinomycin. (b) Whole genome sequencing was performed to verify deletion of *dns*. Sequencing reads map to the *dns* locus for wild-type (top, WT). No reads for this locus are found for the knockout (bottom, KO), confirming deletion of the dns gene.

Taken together, these data demonstrate SXT-mediated recombineering in *V. natriegens*, which can be used to perform episomal DNA manipulations or genomic knock-outs. Efforts in process and host engineering to improve these efficiencies are currently ongoing. For example, identification and inactivation of *E. coli* exonucleases has been shown to improve the *in vivo* stability of both oligonucleotides and double-stranded cassettes, resulting in significant improvement for selection-based recombineering (*33*). Replacing native regulatory elements for SXT may also be beneficial, as SXT-Beta has be found to be under strong translational regulation in *V. cholerae* (*34*). *Finally, CRISPR/Cas9 methods, previously demonstrated in V. natriegens*, can be used as powerful negative selection to enable facile isolation of recombinants (*1*).

Enabling massively-multiplexed oligonucleotide recombineering in *V. natriegens* would allow for the rapid generation of biotechnologically useful strains for cloning, protein expression, and production of valuable molecules (*2–5, 35*). Such strains could be further engineered for virus resistance, incorporation of nonstandard amino acids, and genetic isolation (*4, 36, 37*). Importantly, these methods will also facilitate high-throughput functional genomics for *V. natriegens*, accelerating our ability to study the genetic determinants underlying its remarkable generation time.

## Acknowledgements

This work was supported by Department of Energy Grant DE-FG02-02ER63445.

## Methods

### Growth Media and Chemical concentrations

Standardized growth media for *V. natriegens* is named LB3 - Lysogeny Broth with 3% (w/v) final NaCl. We prepare this media by adding 20 grams of NaCl to 25 grams of LB Broth - Miller (Fisher BP9723-500). All strains were cultured at 37°C. Minimal M9 media was prepared according to manufacturer's instruction with sodium chloride added to 2% (w/v) final. Glucose was added to 0.4% (v/v) final. For M9-FOA media, 1 mg/ml FOA (EZSolution, BioVision) and 20 μg/ml uracil were added to M9. The following concentrations of chemicals were used: Ampicillin/Carbenicillin - 100 μg/ml, Kanamycin - 75 μg/ml, Chloramphenicol - 5μg/ml, Spectinomycin - 100μg/ml, IPTG - 10mM, 5-FOA - 1 mg/ml.

### Strains and Plasmids

All plasmids were transformed by electroporation into *Vibrio natriegens* ATCC 14048. The plasmids in this study are: pRSF-pLac-SXT (AmpR) carrying SXT-Beta, SXT-Exo, and λ-Gam; pRSF-ara-λRed (AmpR) carrying λ-Beta, λ-Exo, and λ-Gam; pRST-brokenspec (KanR) carrying an inactive spectinomycin cassette.

### Electroporation of *V. natriegens*

Electrocompetent *V. natriegens* were prepared as previously described (*1, 38*). Briefly, *V. natriegens* was grown overnight in LB3 at 37°`C`. A subculture was prepared by inoculating an overnight culture, washed once in fresh LB3, at 1:100 dilution into fresh media until OD_600_ ~0.4. Cells were then washed by resuspension in 1ml of cold 1M sorbitol and resuspended in 1M sorbitol. 50μL of concentrated cells were used per transformation. To transform, the indicated amount of DNA was added to the cells in 0.1mm cuvettes and electroporated using Bio-Rad Gene Pulser electroporator at 0.4kV, 1kΩ, 25μF. Cells were recovered in 1mL, unless otherwise indicated, LB3 or SOC3 media for 45 minutes at 37°`C` at 225rpm, and plated on LB3 + 1.5% (w/v) agar with antibiotics as needed. Plates were incubated > 6 hours at 37°`C`.

### Synthesis and cloning of SXT recombination proteins and λ-Red

SXT-Beta and SXT-Exo (GenBank AY055428.1), including the native intergenic regions, were synthesized (Invitrogen GeneArt) from the SXT mobile element and cloned in frame of λ-Beta and λ-Exo using NEBuilder HiFi (NEB) onto the *E. coli* lac-inducible pRSF backbone (*20*). The *E. coli* arabinose-inducible λ-Red operon was cloned from pKD46 (Yale CGSC) onto the pRSF backbone since *V. natriegens* does not replicate the temperature sensitive allele repA101ts on pKD46 at 25°`C` or 30°`C`. It is likely that SXT proteins are more compatible with *Vibrio* hosts compared to λ, which has been evolutionarily optimized for activity with *E. coli*. Furthermore, one previous study demonstrated that λ-Beta could enable dsDNA recombination in *V. cholerae*, though it was 100-fold less efficient and required longer homology arms than in *E. coli* (*15*).

### Spectinomycin selection assay for oligonucleotide-mediated recombination

A spectinomycin gene was inserted (NEBuilder HiFi) onto pRST and a premature stop codon (T431A) was introduced by PCR mutagenesis. This site was chosen since allelic replacement of T:T (chromosomal:synthetic) has been found to be efficient in *E. coli* mutS^+^ strains (*28*). The resulting plasmid is named pRST-brokenspec. For routine experiments, strains were revived from -80°`C` in 3mL of LB3 strain and grown overnight with the appropriate chemicals at 37°`C` in LB3 at 225rpm. Cultures were subcultured in the appropriate media and harvested (OD_600_ ~0.4) for electroporation with oligonucleotides. Following recovery for 45 minutes at 37°`C` at 225rpm, cultures were plated on LB3 plates supplemented with spectinomycin. 90-mer ssDNA oligonucleotides were used as previous studies have shown that recombination efficiency in *E. coli* is optimal with 90-mers (*5, 39*). All oligonucleotides were purchased from IDT.

### FOA selection

*V. natriegens* cells carrying pRST-pLac-SXT plasmid were inoculated from -80°`C` and grown overnight in 3mL supplemented with carbenicillin and IPTG at 37°`C` in LB3 at 225rpm. Cultures were subcultured in the same media and harvested (OD_600_ ~0.4) for electroporation. Oligonucleotides were designed to introduce a premature stop codon (T335A) in *pyrF* (pyrFv1_F, pyrFv1_R; 750 pmol). Following 45 min recovery at 37°`C` in LB3, cells were washed in M9 media twice, plated on M9 plates and incubated at 37°`C` for one hour. After this short incubation, we performed replica plating onto M9-FOA plates. Resulting colonies were PCR amplified and sequenced by Sanger (Genewiz).

### Deep sequencing of oligonucleotide-mediated genome recombination

The corresponding mutations for each target were as follows: *dns*: CTGACT (355) TAATAA; *mutS*: CCTCCT (1240) TAGTAG; lacZ: ACTATA 592 TAATAG; *xds*: GCCGCA 1052 TAATAG. A 10μL mixture of “forward” or “reverse” oligonucleotides, totaling 2500 pmol, were electroporated and cells recovered in 30mL of LB3 at 37°`C` until OD_600_ ~0.4. A second round of recombineering was performed by harvesting these samples for electroporation. Genomic DNA was extracted (GE illustra bacteria genomicPrep) for each sample at OD_600_ ~0.4. Each of the four genomic loci were PCR amplified with Illumina-tailed primers 30 bp upstream and downstream of the targeting 90-mer to ensure specific amplification of the target loci. qPCR was utilized to ensure that all four targets were evenly amplified. The same concentration of genomic DNA was used in each PCR cycle. All targets amplified with the same number of PCR cycles, except for *xds*, which required 2 extra cycles (~4-fold difference in copy number). Samples were barcoded and sequenced with MiSeq v3 150. Sequences were trimmed with cutadapt and the analyzed for evidence of allelic exchange (*40*).

### Whole genome sequencing of *dns* knock-out strain

Genomic DNA (GE illustra bacteria genomicPrep) was harvested from overnight culture of this strain at 37°`C` in LB3. Sequencing libraries were prepared (Illumina Nextera) and sequenced on MiSeq v3 150. Adapters were trimmed with cutadapt and aligned with bowtie to the reference genome, RefSeq NZ_CP009977-8 (*40, 41*).

**Table 1.**
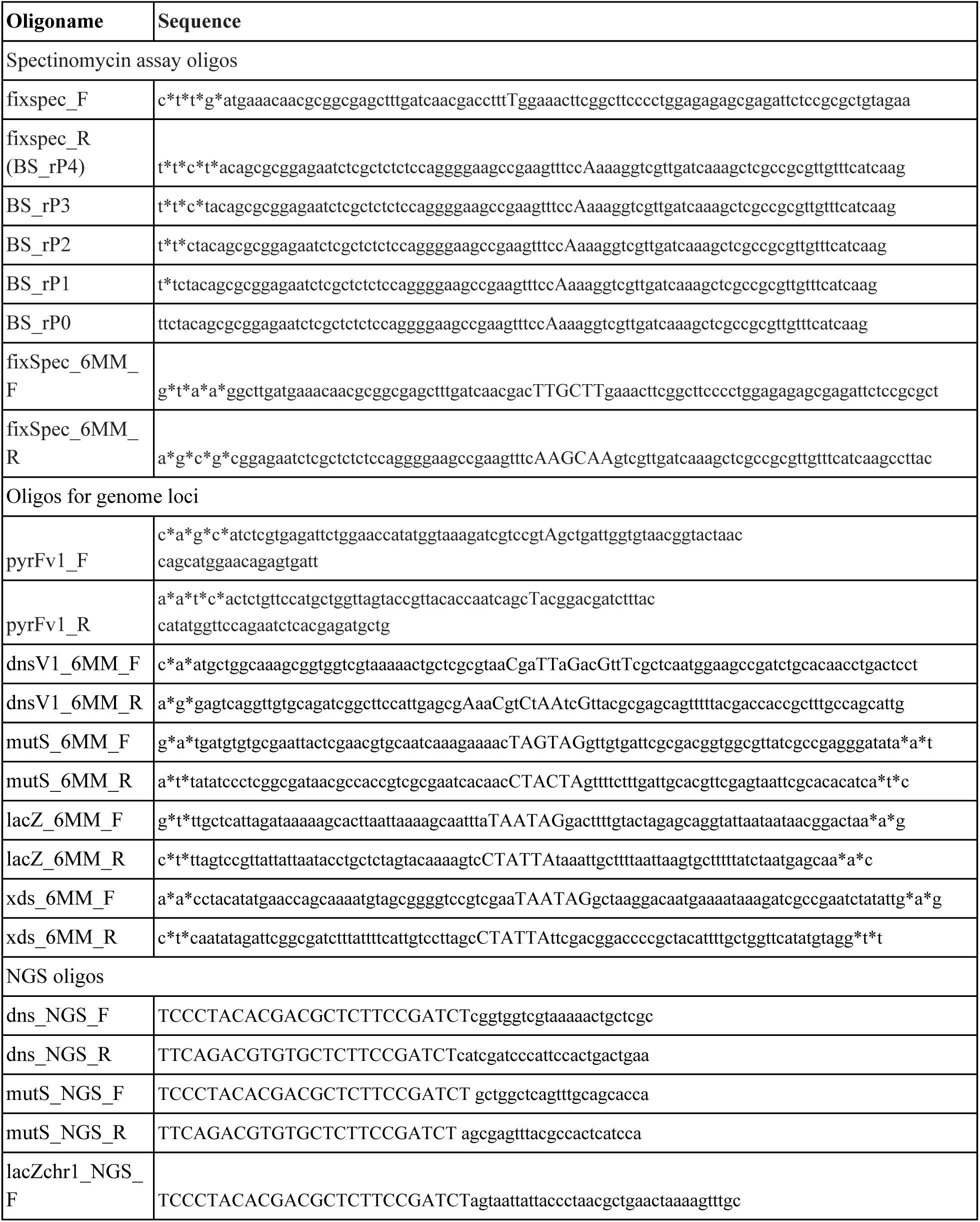
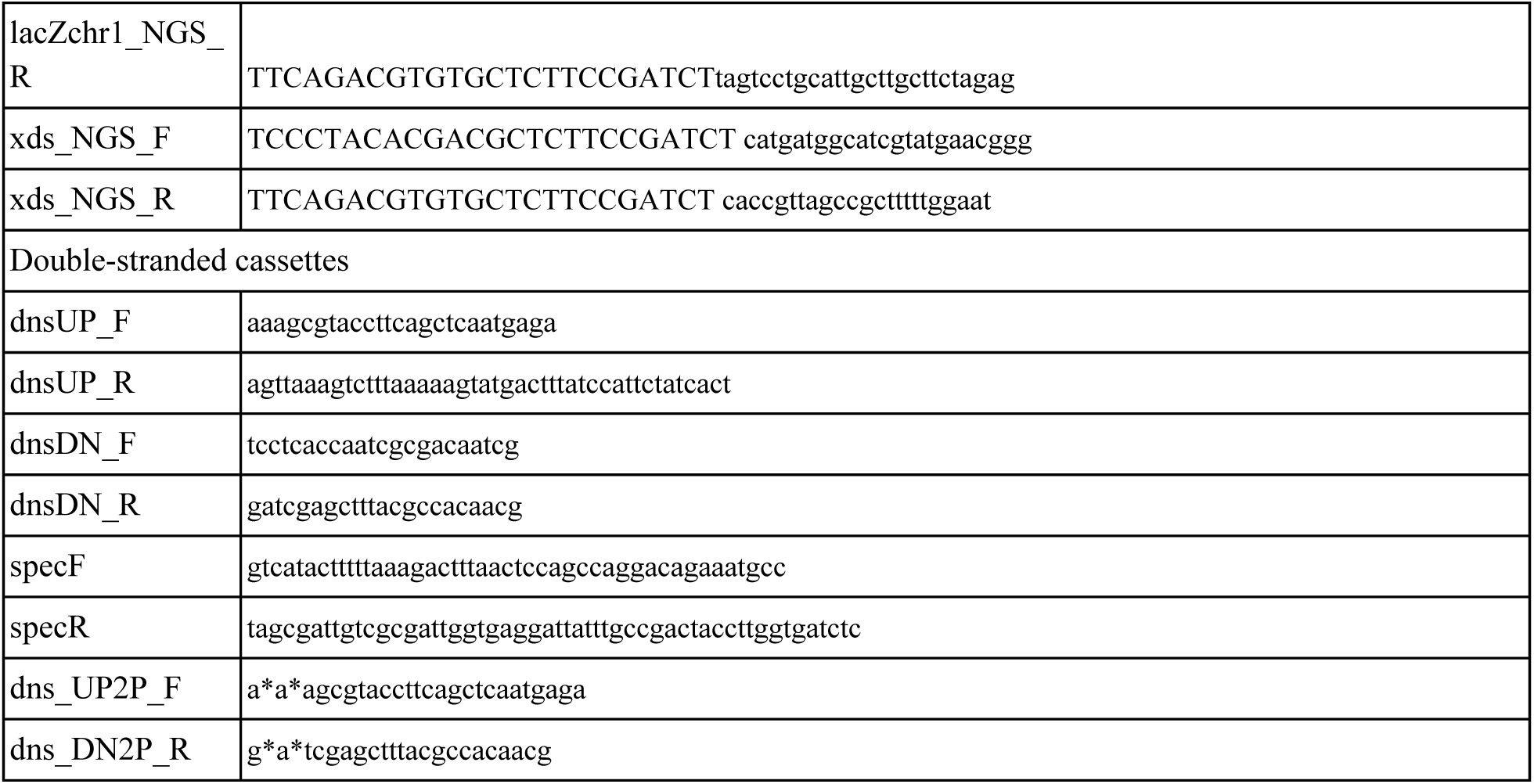
Oligos used in this study.

